# Limited potexvirus diversity in eastern Gulf of Mexico seagrass meadows

**DOI:** 10.1101/2023.12.11.571111

**Authors:** Shen Jean Lim, Karyna Rosario, Meredith E. Kernbach, Anthony J. Gross, Bradley T. Furman, Mya Breitbart

## Abstract

2.

*Turtlegrass virus X*, which infects the seagrass *Thalassia testudinum*, is the only potexvirus known to infect marine flowering plants. We investigated potexvirus distribution in seagrasses using a degenerate reverse transcription polymerase chain reaction (RT-PCR) assay originally designed to capture potexvirus diversity in terrestrial plants. The assay, which implements Potex-5 and Potex-2RC primers, successfully amplified a 584 nt RNA-dependent RNA polymerase (RdRp) fragment from TVX-infected seagrasses. Following validation, we screened 74 opportunistically collected, apparently healthy seagrass samples for potexviruses using this RT-PCR assay. The survey examined the host species *T. testudinum, Halodule wrightii, Halophila stipulacea, Syringodium filiforme, Ruppia maritima*, and *Zostera marina*. Potexvirus PCR products were successfully generated only from *T. testudinum* samples and phylogenetic analysis of sequenced PCR products revealed five distinct TVX sequence variants. Although the RT-PCR assay revealed limited potexvirus diversity in seagrasses, the expanded geographic distribution of TVX shown here emphasizes the importance of future studies to investigate *T. testudinum* populations across its native range and understand how the observed fine-scale genetic diversity a?ects host-virus interactions.

**Impact statement:** Potexviruses are widespread in terrestrial plants; however, the recent discovery of TVX in the seagrass *Thalassia testudinum* extends their host range to marine flowering plants. Here we use existing Potex-5 and Potex-2RC degenerate primers to explore potexvirus infections in several seagrass species. TVX sequence variants were detected in *T. testudinum* collected from the eastern Gulf of Mexico, uncovering previously unknown genetic diversity of this poorly understood virus.

**Data summary:** All sequence data are available in NCBI GenBank under the accession numbers OR827692-OR827705, OR854648, OR863396, OR879052-OR879056, and PP430548-PP430571. The authors confirm all supporting data, code and protocols have been provided within the article or through supplementary data files.

## 5. Introduction

Potexviruses are positive-sense single-stranded RNA (ssRNA) viruses from the family *Alphaflexiviridae* that infect many agronomically important flowering plants (angiosperms), including potato, tobacco, and tomato (1). The type member of this genus is *Potato virus X* (PVX), from which the genus name *Potexvirus* is derived (2). Potexviruses have a broad geographic range that reflects the distribution of their hosts and the prevalence of global commerce (2, 3). Although each potexvirus species has a limited natural host range, some species can infect a wide range of host plants in experimental settings (1). Potexviruses are transmitted mechanically through contaminated agricultural equipment or plant-to-plant contact (1-3). Transmission via seed and aphid vectors and non-specific transmission through chewing insects has been reported but is less common (1-3). Most potexviruses cause persistent, asymptomatic infections or mild mosaic disease, while some cause dwarf, necrotic, or ringspot symptoms in their natural hosts (1, 2). Notably, PVX can cause severe symptoms when its plant host is co-infected with potyviruses (4-6).

In addition to infecting terrestrial plants, potexviruses are also capable of infecting seagrasses, the only marine angiosperms. A novel potexvirus species, *Turtlegrass virus X* (TVX), was discovered in apparently healthy *Thalassia testudinum* seagrass samples collected from Terra Ceia Aquatic Preserve, Tampa Bay, Florida, USA (7). The identification of TVX indicates that an important aspect of potexvirus diversity has been overlooked by focusing only on terrestrial plants. Degenerate primers used in reverse transcriptase polymerase chain reaction (RT-PCR) assays, such as Potex-5 and Potex-2RC (8), have proven useful in detecting diverse potexviruses, but have not been tested in aquatic habitats. This study applied the Potex-5 and Potex-2RC primer pair in a RT-PCR assay to explore potexvirus diversity in samples from six seagrass host species.

## 6. Methods

### Primer validation

Potex-5/Potex-2RC were tested on cDNA samples each synthesized from 1 μg RNA extracted from *Portulaca* sp. infected with *Alternanthera mosaic virus* (AltMV) and *Opuntia* sp. with a mixed infection of potexviruses, as well as cDNA pooled from *T. testudinum* samples known to contain TVX, based on the TVX-specific RT-PCR assay described by Van Bogaert *et al*. 2019 (7). The PCR reaction contained 0.48 μM of each primer, 2 μL cDNA template, 1 μL GC enhancer, and 1X AmpliTaq Gold™ 360 Master Mix (Applied Biosystems™, Waltham, MA, USA) in a 25 μL reaction volume. PCR was performed under the following conditions: initial denaturation at 95°C for 10 minutes, 40 cycles of denaturation at 95°C for 30 seconds, annealing at 51.5°C (as published in van der Vlugt and Berendsen (8)) for 30 seconds, extension at 72°C for 1 minute, followed by elongation at 72°C for 10 minutes and cooling at 11°C. The PCR product was visualized following gel electrophoresis on a 1% (wt/vol) agarose gel stained with ethidium bromide. All PCR reactions, except for the no template control, yielded visible bands. All PCR products were purified using the Zymoclean Gel DNA Recovery Kit (Irvine, CA, USA), quantified using the Qubit^TM^ DNA high sensitivity (HS) assay (Invitrogen™, Waltham, MA, USA), and Sanger sequenced bidirectionally by Eurofins Genomics (Louisville, KY, USA).

### RT-PCR

Total RNA extraction was performed on 30-100 mg of leaves from multiple shoots pooled by seagrass species and collection site (**Table S1**) using Zymo Research’s (ZR) Quick-RNA™ Plant Miniprep kit. Each pooled seagrass sample was homogenized in a BashingBead Lysis Tube containing 2 mm ceramic beads and 800 μL RNA lysis bu?er (provided in the kit) for 5 minutes at maximum speed using a Fisherbrand^TM^ Bead Mill 4 Homogenizer (Fisher Scientific, Waltham, MA, USA). Tissue homogenates were centrifuged at maximum speed (21,130 *g*) for 1 minute and RNA was extracted from the total volume (∼800 μL) of the supernatant. To ensure successful RNA extraction, after each round of extraction, a random subset of RNA samples was quantified using the Qubit^TM^ RNA high sensitivity (HS) assay (Invitrogen™). From each sample, cDNA was synthesized from 8 μL RNA using the SuperScript^TM^ IV First-Strand Synthesis System (Invitrogen™) and following manufacturer’s instructions for random hexamers. PCR amplification was performed on each cDNA sample, followed by gel electrophoresis, PCR product purification, DNA quantification, and Sanger sequencing, using the methods described above.

### Amplicon sequence analysis

Amplicon sequences were first compared with sequences from NCBI’s nucleotide (nt) and non-redundant protein (nr) collections (9) and the TVX genome (7) using the megablast and/or blastx programs on NCBI’s BLAST® server (10). Sequence reads from each sample were then mapped to the Potex-5/Potex-2RC amplicon region (with forward and reverse primer sequences removed) in the TVX genome (7) using the “Map Sanger Reads to Reference” function implemented in Unipro UGENE v48.1 (11), with a trimming quality threshold of 20 and mapping minimum similarity of 70%. Electropherograms of mapped reads were manually inspected to check for mixed infection of potexviruses, to remove primer sequences and low-quality bases, and to resolve ambiguous bases.

Sequences from 42 Potex-5/Potex-2RC PCR products amplified from seagrass and terrestrial plant samples described above were aligned with corresponding amplicon regions (excluding primer sequences) extracted from the reference genomes of TVX (NC_040644) (7), *Bamboo mosaic virus* (NC_001642) (12), *Foxtail mosaic virus* (NC_001483) (13), AltMV (OR607766), PVX (NC_011620) (14), and *Lolium latent virus* (NC_010434) (15) retrieved from the NCBI Virus database (16). Multiple sequence alignment was performed using the L-INS-I and --adjustdirectionaccurately options in MAFFT v7.5.08 (17), then trimmed to 514 nt to remove leading and trailing gaps. Pairwise sequence identities were calculated from the trimmed alignment using Clustal Omega v1.2.3 (18) with the --distmat-out, --percent-id, and –full options. For phylogenetic analysis, Molecular Evolutionary Genetics Analysis (MEGA) 11 (19) identified the best model for the trimmed alignment to be the Hasegawa-Kishino-Yano model (20) with a discrete Gamma distribution (HKY+G). Using this substitution model, a maximum likelihood tree with 1,000 bootstrap replicates was constructed from the alignment using MEGA11 (19). The tree was condensed using MEGA11 (19) to eliminate interior branches with <50% bootstrap support. The final tree was visualized together with the heatmap of pairwise sequence identity using the ggtree R package v3.6.2 (21).

## 7. Results

The Potex-5/Potex-2RC primer pair successfully amplified TVX from *T. testudinum* samples collected from Terra Ceia Aquatic Preserve, AltMV infecting *Portulaca* sp., and an uncharacterized potexvirus infecting *Opuntia* sp. (8). A RT-PCR assay using Potex-5/Potex-2RC was subsequently performed on seagrass leaves opportunistically collected from separate sampling efforts, including 1) *T. testudinum, H. wrightii, S. filiforme*, and *R. maritima* from Tampa Bay on Jan 26^th^ or Feb 4^th^, 2022; 2) *T. testudinum* from the Dry Tortugas National Park, Florida, USA collected between May 16^th^-20^th^, 2022; 3) *T. testudinum* from a systematic seagrass survey at Terra Ceia Aquatic Preserve on August 1^st^, 2022; and 4) *T. testudinum, H. wrightii*, and *S. filiforme* samples collected from Tampa Bay seagrass site S3T8 (Lassing Park) on October 3^rd^, 2023 (**Figure 1** and **Table S1**). Seagrass samples (**Table S1**) were also received from other research groups, including *T. testudinum* from Panacea (Florida, USA), *Zostera marina* from York River (Virginia, USA), West Falmouth Harbor (Massachusetts, USA), Sitka (Alaska, USA), New Zealand, and Kalmar (Sweden), and *Halophila stipulacea* from Jobos Bay National Estuarine Research Reserve (Puerto Rico, USA). PCR products were successfully amplified and sequenced from 40 *T. testudinum* samples from Florida sites, including Terra Ceia Aquatic Preserve, Tampa Bay seagrass sites S1T5 and S3T8 (Lassing Park), Panacea located in the Florida Panhandle, and Florida Keys sites including Bush Key, Garden Key, Marquesas Key, and Key West (**Figure 1**). Based on our analysis of Sanger sequencing traces, no evidence of mixed potexvirus infection was found in these *T. testudinum* samples. Aside from *T. testudinum*, no other seagrass species produced PCR products.

**Figure 1.**
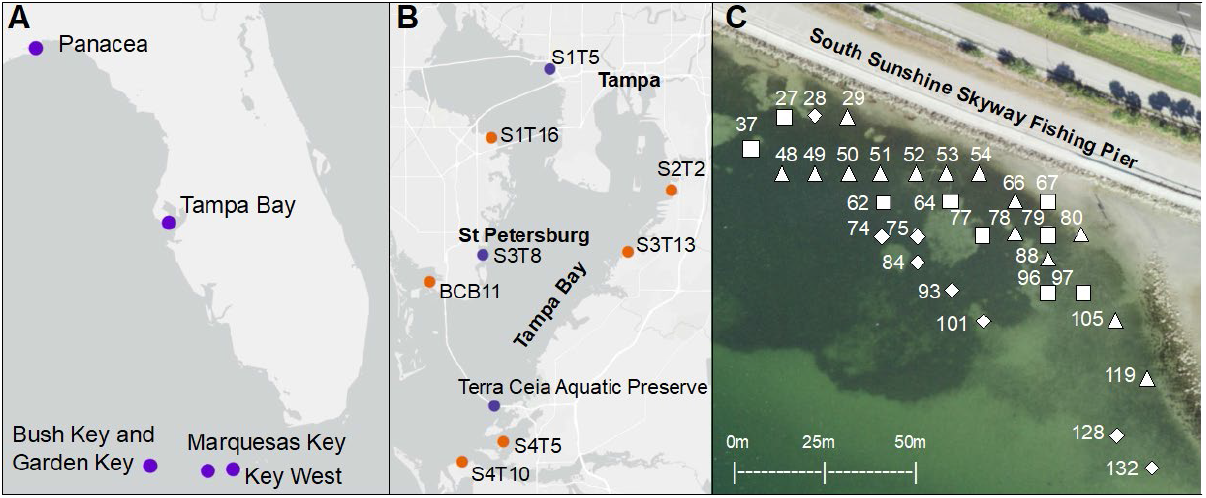
Map showing locations of seagrass sites in (A) Florida, (B) Tampa Bay, and (C) Terra Ceia Aquatic Preserve where samples were collected for RT-PCR. Purple markers in (A) and (B) denote sites where TVX products were amplified from *T. testudinum* with the Potex-5/Potex-2RC primers, whereas samples from sites with orange markers tested negative. Marker shapes in (C) represent potexvirus subclades found in Terra Ceia Aquatic Preserve sampling sites and correspond to **Figure 2** (negative samples not shown). Metadata of seagrass samples is described in **Table S1**.

Phylogenetic analysis placed all Potex-5/Potex-2RC amplicons from seagrasses in the same clade as the RdRp sequence fragment from the TVX genome (7) with 100% bootstrap confidence (**Figure 2A**). Sequences within this clade shared 74.4% to 98.6% nucleotide sequence identities to each other (**Figure 2B**). TVX sequences from Terra Ceia Aquatic Preserve formed three subclades. Average pairwise nucleotide sequence identities were 95.8±1.4% within each subclade and 79.3±2.9% between subclades (**Figure 2**). Two Terra Ceia Aquatic Preserve subclades, including the subclade containing the amplicon region from the TVX genome, were sister groups to each other, sharing 84.6±1% average pairwise nucleotide sequence identities (**Figure 2**). Other TVX sequences from Key West, Marquesas Key, and Panacea formed a subclade with 99.6±0.2% average pairwise nucleotide sequence identities, while sequences from Garden Key and Bush Key formed another subclade with 89.3±0.9% average pairwise nucleotide sequence identities (**Figure 2**). Potexviruses that infect terrestrial grasses (*Foxtail mosaic virus* and *Bamboo mosaic virus*) formed sister clades to the TVX-containing clade (**Figure 2A**).

**Figure 2.**
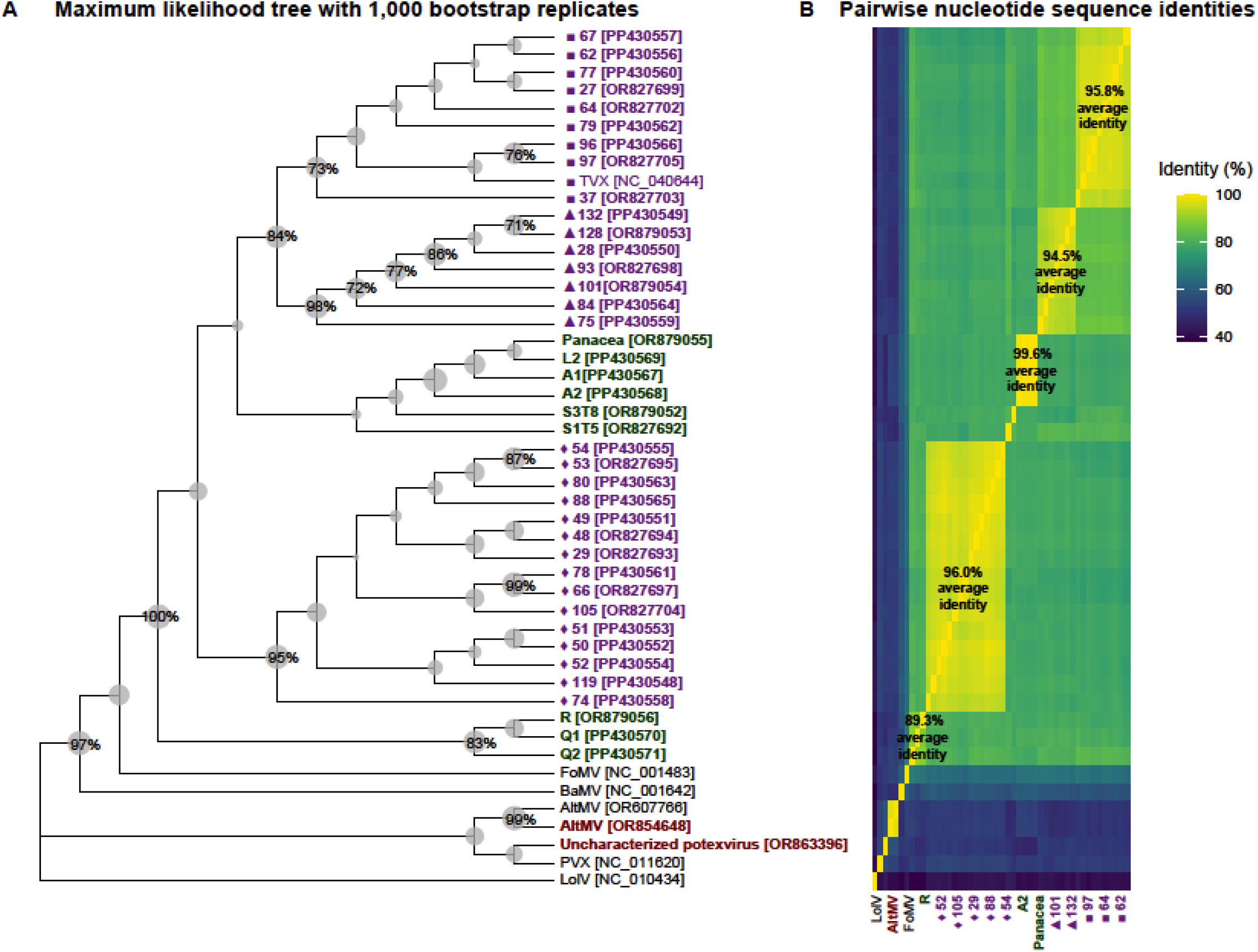
(A) Bootstrap maximum likelihood tree and (B) heatmap of pairwise nucleotide sequence identities of TVX nucleotide sequences amplified from seagrasses collected from Terra Ceia Aquatic Preserve (purple text) and other sites in the eastern Gulf of Mexico (green text), in relation to potexvirus sequences amplified from terrestrial plants (brown text) and potexvirus RdRp sequences from reference genomes (black text). The NCBI accession number for each sequence in (A) is indicated in square brackets. The tree in (A) was constructed by MEGA11 from a 514 nt conserved region in the multiple sequence alignment. Sizes of tree nodes are proportionate to bootstrap values indicating the percentage of trees, based on 1,000 replicates, in which sequences within a node were clustered together. Only bootstrap values >70% are labelled on the tree nodes. *Lolium latent virus* was used as an outgroup for the tree. Marker shapes in (A) and (B) preceding sequences in the Terra Ceia Aquatic Preserve subclades correspond to marker shapes in **Figure 1C**. Abbreviations: TVX, *Turtlegrass virus X*; AltMV, *Alternanthera mosaic virus*; PVX, *Potato virus X*; LolV, *Lolium latent virus*; FoMV, *Foxtail mosaic virus*; BaMV, *Bamboo mosaic virus*.

## 8. Discussion

In this study, we validated the ability of the potexvirus degenerate primers, Potex-5/Potex-2RC, published by van der Vlugt and Berendsen (8), to amplify TVX-infected seagrass samples and potexvirus-infected terrestrial plant samples. Potex-5/Potex-2RC were used in a RT-PCR survey to test six seagrass species for potexviruses. Among the seagrass species examined, potexvirus infection was only detected in *T. testudinum*, the species from which TVX was first identified. The criteria published by the International Committee on Taxonomy of Viruses (ICTV) demarcates potexviruses into species based on host range, absence of cross-protection in infected plants, serology, and sequence identity thresholds of 72% for nucleotide sequences and 80% for amino acid sequences (https://ictv.global/report/chapter/alphaflexiviridae/alphaflexiviridae/potexvirus). Potexviruses amplified in this study are specific to *T. testudinum* and shared >72% sequence identity to each other and to the RdRp sequence fragment from the TVX genome (7). Therefore, we considered identified potexviruses the same species as TVX. Note that this study only analyzed a short and conserved fragment of the RdRp sequence and these pairwise identities may change when considering complete genomes. Nevertheless, our results suggest limited potexvirus diversity in marine angiosperms. RT-PCR analysis revealed *T. testudinum* as a natural host for TVX and confirmed TVX infection in eastern Gulf of Mexico seagrass meadows, including those in northwest (Panacea), west-central (Tampa Bay), and southwest (Florida Keys) Florida. Previous phylogenetic analyses of the PVX coat protein and genomic sequences identified phylogroups that reflect intraspecies diversity (22, 23). Here, three distinct TVX phylogroups, sharing >95% intra-group and <80% inter-group amplicon sequence identity, were observed within a single seagrass meadow at Terra Ceia Aquatic Preserve, along with two additional phylogroups in other Tampa Bay sites, namely the Florida Keys and Panacea.

Among all sites sampled in this study, the most extensive TVX survey was conducted within the Terra Ceia Aquatic Preserve. Systematic RT-PCR surveys at other sites will be useful in elucidating potexvirus diversity in seagrasses. The sampling scheme in this study was limited to *T. testudinum* populations along the eastern Gulf of Mexico. However, *T. testudinum* has a widespread geographic range across the Western Atlantic Ocean with high genetic diversity that is generally partitioned into two clusters corresponding to the Gulf of Mexico or the Caribbean phylogeographic regions (24). Although seagrasses can reproduce asexually, some *T. testudinum* populations, such as those in Tampa Bay (25-27) and Florida Bay (28, 29), undergo sexual reproduction and exhibit high genotypic diversity. Currently, the relationship between *T. testudinum* genotype and TVX diversity is unknown and should be further investigated. Future potexvirus surveys covering the total distribution range of *T. testudinum*, the related species *T. hemprichii*, and other seagrass species also have the potential to uncover relationships between potexvirus diversity, host diversity, and their phylogeographic distributions.

## 9. Author statements

### 9.1 Author contributions

SJL: Data curation, Formal analysis, Investigation, Methodology, Software, Supervision, Visualization, Writing - original draft

KRC: Conceptualization, Formal analysis, Funding acquisition, Writing – review & editing. MEK: Investigation, Writing – review & editing

AJG: Investigation, Writing - review & editing

BTF: Funding acquisition, Investigation, Writing - review & editing

MB: Conceptualization, Formal analysis, Funding acquisition, Investigation, Methodology, Project administration, Resources, Supervision, Validation, Writing - review & editing.

### 9.4 Ethical approval

N/A

### 9.5 Consent for publication

N/A

## 9.6 Acknowledgements

Fieldwork was conducted under permit no. FKNMS-2022-026 issued by the Florida Keys National Marine Sanctuary, permit no. DRTO-2022-SCI-0004 issued through the U.S. Department of the Interior National Park Service and permit no. SAL-19-1172A-SR issued through the Florida Fish and Wildlife Conservation Commission. We acknowledge the contributions of Luis Alvarado-Marchena, Victoria Congdon, Keith Keel, Isabella Ritchie, Taylor Little, Jessye Kirkham, and Mike Wheeler in field work and sample collection at these sites. Isabella Ritchie provided cDNAs from *T. testudinum* samples from Terra Ceia Aquatic Preserve. Salvador P Lopez, Jr and Scott Adkins from the Agricultural Research Service, United States Horticultural Research Laboratory, U.S. Department of Agriculture (USDA), Fort Pierce, FL, USA provided cDNAs from potexvirus-infected plant samples. We thank Maya Groner, Ian Hewson, Karin Holmfeldt, Natalia Lopez Figueroa, Carolyn Malmstrom, Jordan Rede, Cypress Rudloe, and Jack Rudloe for providing seagrass samples from other sites.

---

**Table S1.**
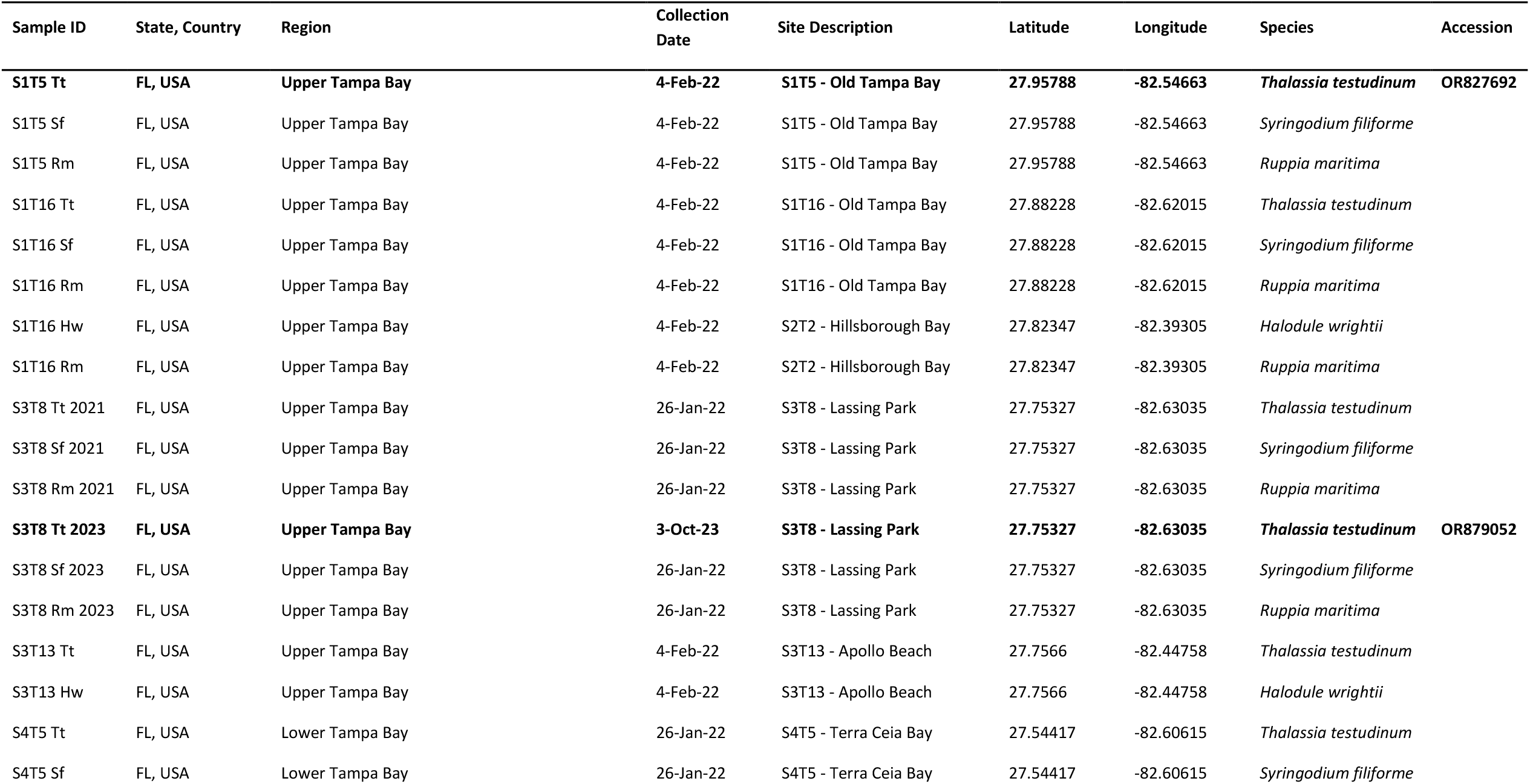
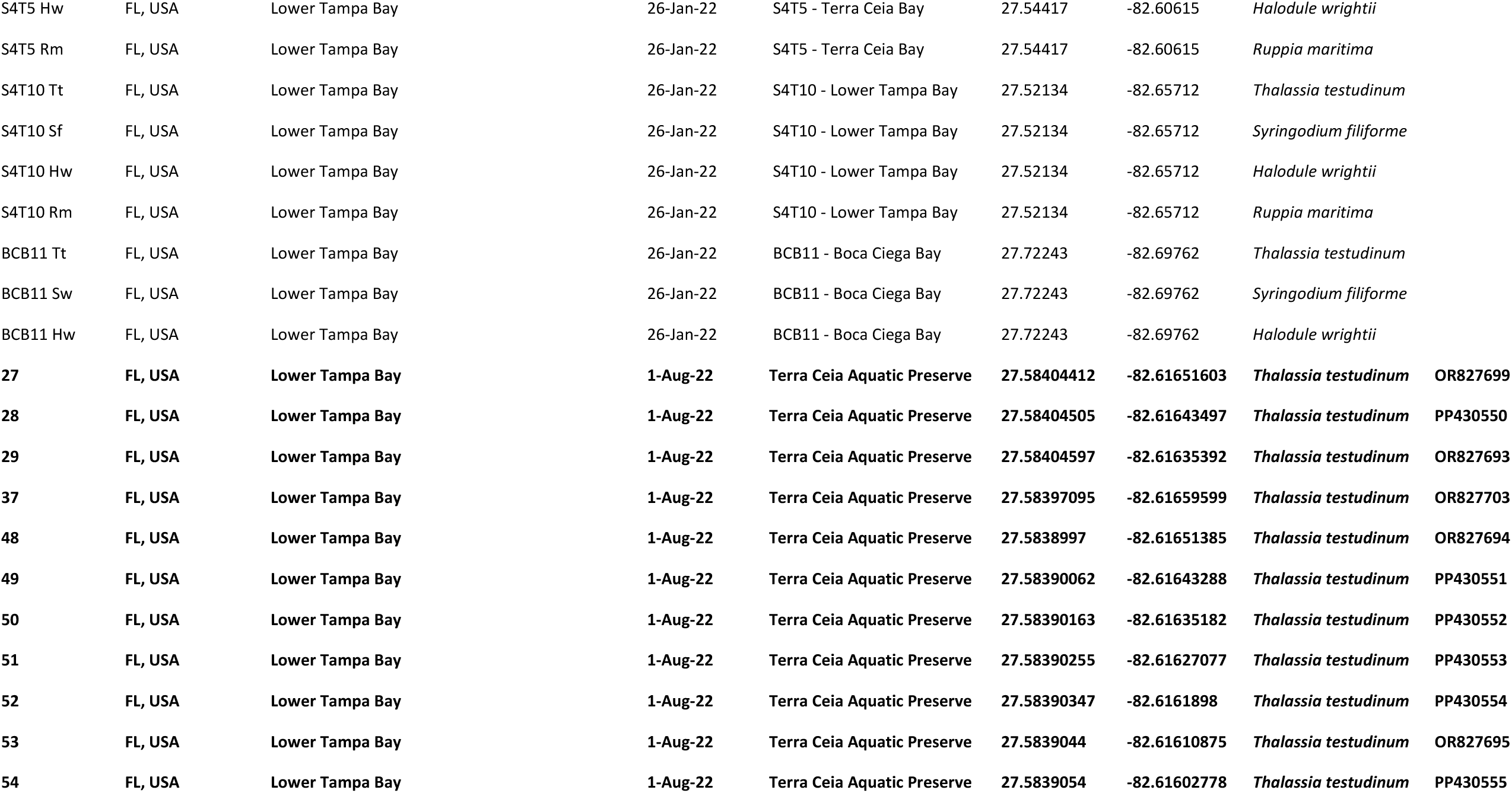
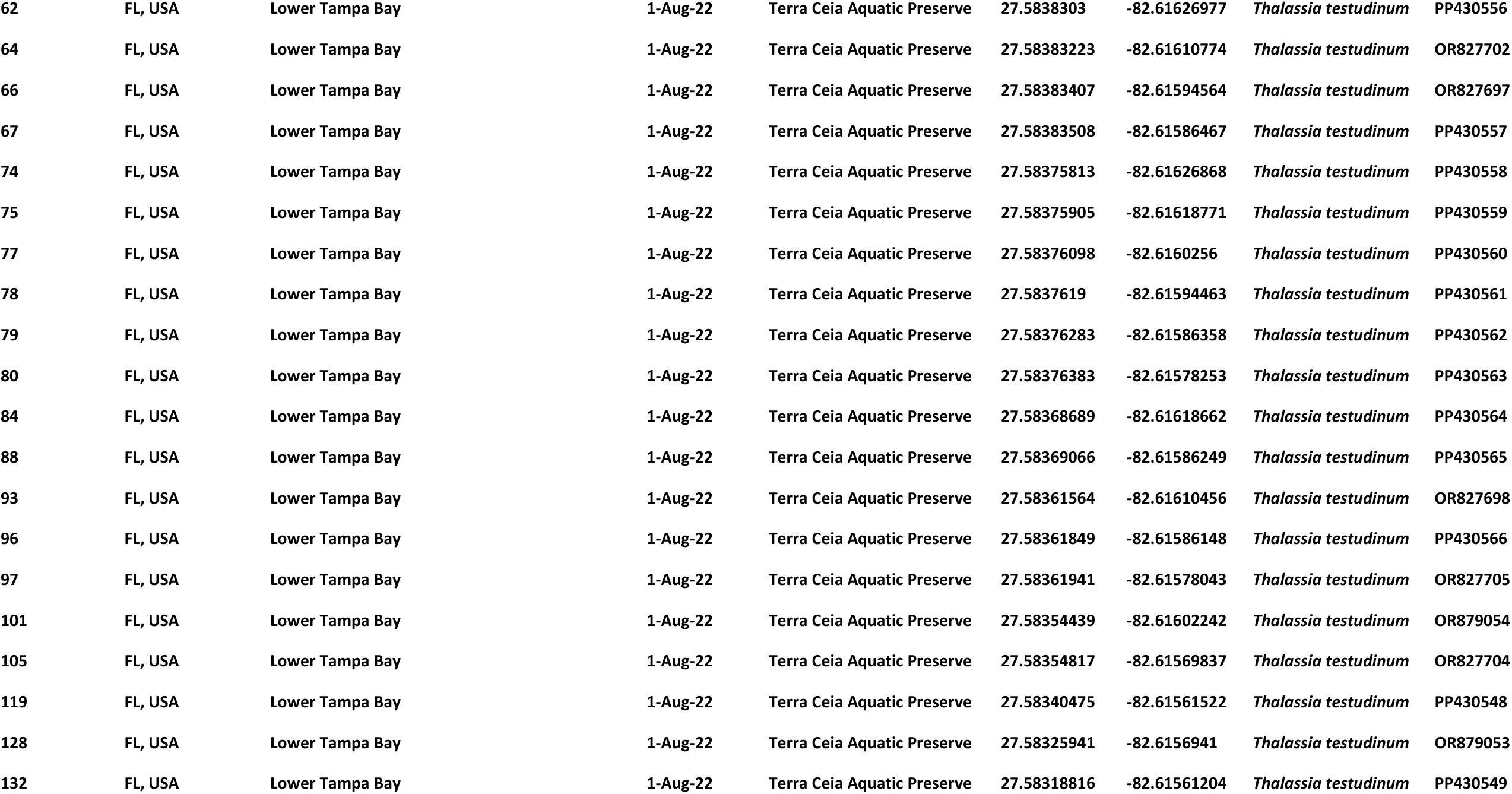
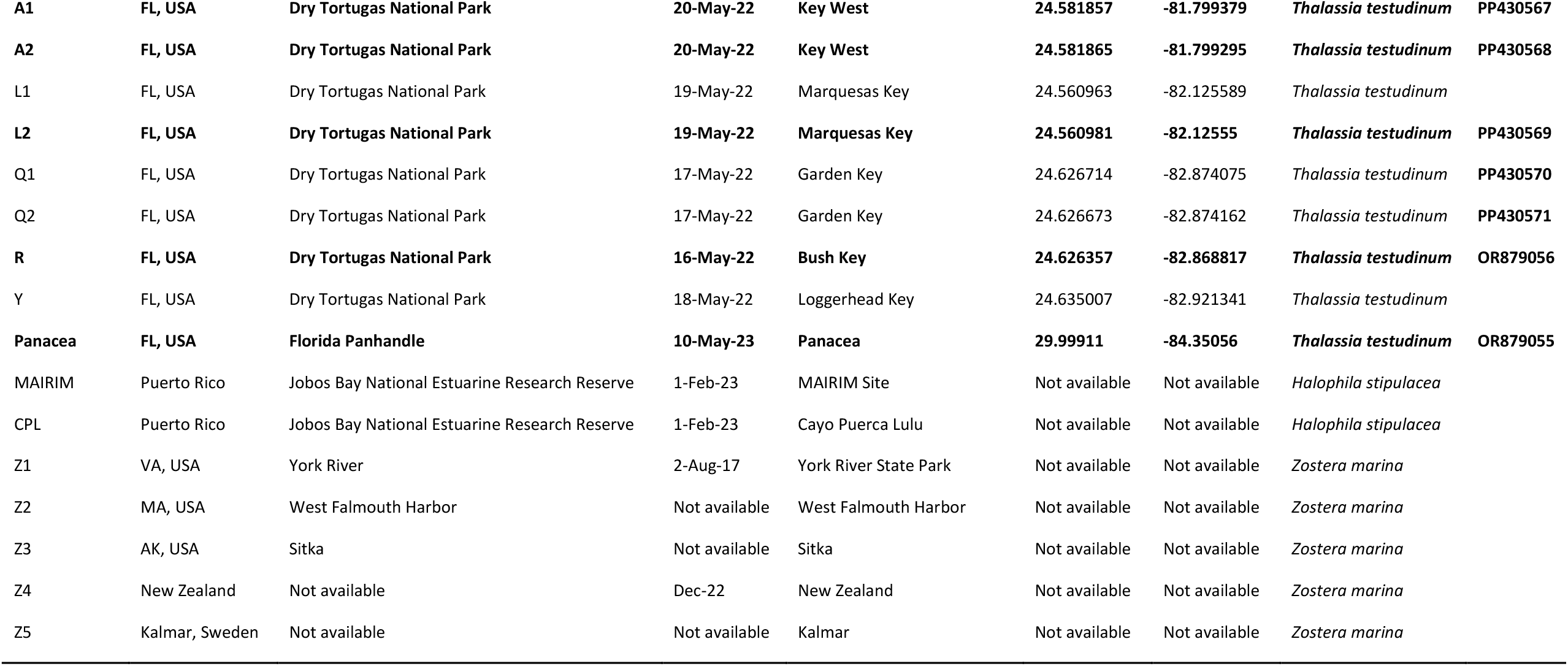
Metadata of seagrass samples used in the RT-PCR survey. Samples successfully amplified by Potex-5/Potex-2RC primers are highlighted in bold and the NCBI accession numbers of their amplicon sequences are listed in the last column.

