## Supplementary material for "Limited potexvirus diversity in eastern Gulf of Mexico seagrass meadows": Table S1

**Table S1**. Metadata of seagrass samples used in the RT-PCR survey. Samples successfully amplified by Potex5F/Potex2-RC primers are highlighted in bold and the NCBI accession numbers of their amplicon sequences are listed in the last column.

| Sample ID | State, Country | Region | Collection Date | Site Description | Latitude | Longitude | Species | Accession |
|---|---|---|---|---|---|---|---|---|
| **S1T5 Tt** | **FL, USA** | **Upper Tampa Bay** | **4-Feb-22** | **S1T5 - Old Tampa Bay** | **27.95788** | **-82.54663** | ***Thalassia testudinum*** | **OR827692** |
| S1T5 Sf | FL, USA | Upper Tampa Bay | 4-Feb-22 | S1T5 - Old Tampa Bay | 27.95788 | -82.54663 | *Syringodium filiforme* | |
| S1T5 Rm | FL, USA | Upper Tampa Bay | 4-Feb-22 | S1T5 - Old Tampa Bay | 27.95788 | -82.54663 | *Ruppia maritima* | |
| S1T16 Tt | FL, USA | Upper Tampa Bay | 4-Feb-22 | S1T16 - Old Tampa Bay | 27.88228 | -82.62015 | *Thalassia testudinum* | |
| S1T16 Sf | FL, USA | Upper Tampa Bay | 4-Feb-22 | S1T16 - Old Tampa Bay | 27.88228 | -82.62015 | *Syringodium filiforme* | |
| S1T16 Rm | FL, USA | Upper Tampa Bay | 4-Feb-22 | S1T16 - Old Tampa Bay | 27.88228 | -82.62015 | *Ruppia maritima* | |
| S1T16 Hw | FL, USA | Upper Tampa Bay | 4-Feb-22 | S2T2 - Hillsborough Bay | 27.82347 | -82.39305 | *Halodule wrightii* | |
| S1T16 Rm | FL, USA | Upper Tampa Bay | 4-Feb-22 | S2T2 - Hillsborough Bay | 27.82347 | -82.39305 | *Ruppia maritima* | |
| S3T8 Tt 2021 | FL, USA | Upper Tampa Bay | 26-Jan-22 | S3T8 - Lassing Park | 27.75327 | -82.63035 | *Thalassia testudinum* | |
| S3T8 Sf 2021 | FL, USA | Upper Tampa Bay | 26-Jan-22 | S3T8 - Lassing Park | 27.75327 | -82.63035 | *Syringodium filiforme* | |
| S3T8 Rm 2021 | FL, USA | Upper Tampa Bay | 26-Jan-22 | S3T8 - Lassing Park | 27.75327 | -82.63035 | *Ruppia maritima* | |
| **S3T8 Tt 2023** | **FL, USA** | **Upper Tampa Bay** | **3-Oct-23** | **S3T8 - Lassing Park** | **27.75327** | **-82.63035** | ***Thalassia testudinum*** | **OR879052** |
| S3T8 Sf 2023 | FL, USA | Upper Tampa Bay | 26-Jan-22 | S3T8 - Lassing Park | 27.75327 | -82.63035 | *Syringodium filiforme* | |
| S3T8 Rm 2023 | FL, USA | Upper Tampa Bay | 26-Jan-22 | S3T8 - Lassing Park | 27.75327 | -82.63035 | *Ruppia maritima* | |
| S3T13 Tt | FL, USA | Upper Tampa Bay | 4-Feb-22 | S3T13 - Apollo Beach | 27.7566 | -82.44758 | *Thalassia testudinum* | |
| S3T13 Hw | FL, USA | Upper Tampa Bay | 4-Feb-22 | S3T13 - Apollo Beach | 27.7566 | -82.44758 | *Halodule wrightii* | |
| S4T5 Tt | FL, USA | Lower Tampa Bay | 26-Jan-22 | S4T5 - Terra Ceia Bay | 27.54417 | -82.60615 | *Thalassia testudinum* | |
| S4T5 Sf | FL, USA | Lower Tampa Bay | 26-Jan-22 | S4T5 - Terra Ceia Bay | 27.54417 | -82.60615 | *Syringodium filiforme* | |
| S4T5 Hw | FL, USA | Lower Tampa Bay | 26-Jan-22 | S4T5 - Terra Ceia Bay | 27.54417 | -82.60615 | *Halodule wrightii* | |
| S4T5 Rm | FL, USA | Lower Tampa Bay | 26-Jan-22 | S4T5 - Terra Ceia Bay | 27.54417 | -82.60615 | *Ruppia maritima* | |
| S4T10 Tt | FL, USA | Lower Tampa Bay | 26-Jan-22 | S4T10 - Lower Tampa Bay | 27.52134 | -82.65712 | *Thalassia testudinum* | |
| S4T10 Sf | FL, USA | Lower Tampa Bay | 26-Jan-22 | S4T10 - Lower Tampa Bay | 27.52134 | -82.65712 | *Syringodium filiforme* | |
| S4T10 Hw | FL, USA | Lower Tampa Bay | 26-Jan-22 | S4T10 - Lower Tampa Bay | 27.52134 | -82.65712 | *Halodule wrightii* | |
| S4T10 Rm | FL, USA | Lower Tampa Bay | 26-Jan-22 | S4T10 - Lower Tampa Bay | 27.52134 | -82.65712 | *Ruppia maritima* | |
| BCB11 Tt | FL, USA | Lower Tampa Bay | 26-Jan-22 | BCB11 - Boca Ciega Bay | 27.72243 | -82.69762 | *Thalassia testudinum* | |
| BCB11 Sw | FL, USA | Lower Tampa Bay | 26-Jan-22 | BCB11 - Boca Ciega Bay | 27.72243 | -82.69762 | *Syringodium filiforme* | |

| | | | | | | | | | |
|---|---|---|---|---|---|---|---|---|---|
| BCB11 Hw | FL, USA | Lower Tampa Bay | | 26-Jan-22 | BCB11 - Boca Ciega Bay | 27.72243 | -82.69762 | *Halodule wrightii* | |
| **27** | **FL, USA** | **Lower Tampa Bay** | | **1-Aug-22** | **Terra Ceia Aquatic Preserve** | **27.58404412** | **-82.61651603** | ***Thalassia testudinum*** | **OR827699** |
| **28** | **FL, USA** | **Lower Tampa Bay** | | **1-Aug-22** | **Terra Ceia Aquatic Preserve** | **27.58404505** | **-82.61643497** | ***Thalassia testudinum*** | **PP430550** |
| **29** | **FL, USA** | **Lower Tampa Bay** | | **1-Aug-22** | **Terra Ceia Aquatic Preserve** | **27.58404597** | **-82.61635392** | ***Thalassia testudinum*** | **OR827693** |
| **37** | **FL, USA** | **Lower Tampa Bay** | | **1-Aug-22** | **Terra Ceia Aquatic Preserve** | **27.58397095** | **-82.61659599** | ***Thalassia testudinum*** | **OR827703** |
| **48** | **FL, USA** | **Lower Tampa Bay** | | **1-Aug-22** | **Terra Ceia Aquatic Preserve** | **27.5838997** | **-82.61651385** | ***Thalassia testudinum*** | **OR827694** |
| **49** | **FL, USA** | **Lower Tampa Bay** | | **1-Aug-22** | **Terra Ceia Aquatic Preserve** | **27.58390062** | **-82.61643288** | ***Thalassia testudinum*** | **PP430551** |
| **50** | **FL, USA** | **Lower Tampa Bay** | | **1-Aug-22** | **Terra Ceia Aquatic Preserve** | **27.58390163** | **-82.61635182** | ***Thalassia testudinum*** | **PP430552** |
| **51** | **FL, USA** | **Lower Tampa Bay** | | **1-Aug-22** | **Terra Ceia Aquatic Preserve** | **27.58390255** | **-82.61627077** | ***Thalassia testudinum*** | **PP430553** |
| **52** | **FL, USA** | **Lower Tampa Bay** | | **1-Aug-22** | **Terra Ceia Aquatic Preserve** | **27.58390347** | **-82.6161898** | ***Thalassia testudinum*** | **PP430554** |
| **53** | **FL, USA** | **Lower Tampa Bay** | | **1-Aug-22** | **Terra Ceia Aquatic Preserve** | **27.5839044** | **-82.61610875** | ***Thalassia testudinum*** | **OR827695** |
| **54** | **FL, USA** | **Lower Tampa Bay** | | **1-Aug-22** | **Terra Ceia Aquatic Preserve** | **27.5839054** | **-82.61602778** | ***Thalassia testudinum*** | **PP430555** |
| **62** | **FL, USA** | **Lower Tampa Bay** | | **1-Aug-22** | **Terra Ceia Aquatic Preserve** | **27.5838303** | **-82.61626977** | ***Thalassia testudinum*** | **PP430556** |
| **64** | **FL, USA** | **Lower Tampa Bay** | | **1-Aug-22** | **Terra Ceia Aquatic Preserve** | **27.58383223** | **-82.61610774** | ***Thalassia testudinum*** | **OR827702** |
| **66** | **FL, USA** | **Lower Tampa Bay** | | **1-Aug-22** | **Terra Ceia Aquatic Preserve** | **27.58383407** | **-82.61594564** | ***Thalassia testudinum*** | **OR827697** |
| **67** | **FL, USA** | **Lower Tampa Bay** | | **1-Aug-22** | **Terra Ceia Aquatic Preserve** | **27.58383508** | **-82.61586467** | ***Thalassia testudinum*** | **PP430557** |
| **74** | **FL, USA** | **Lower Tampa Bay** | | **1-Aug-22** | **Terra Ceia Aquatic Preserve** | **27.58375813** | **-82.61626868** | ***Thalassia testudinum*** | **PP430558** |
| **75** | **FL, USA** | **Lower Tampa Bay** | | **1-Aug-22** | **Terra Ceia Aquatic Preserve** | **27.58375905** | **-82.61618771** | ***Thalassia testudinum*** | **PP430559** |
| **77** | **FL, USA** | **Lower Tampa Bay** | | **1-Aug-22** | **Terra Ceia Aquatic Preserve** | **27.58376098** | **-82.6160256** | ***Thalassia testudinum*** | **PP430560** |
| **78** | **FL, USA** | **Lower Tampa Bay** | | **1-Aug-22** | **Terra Ceia Aquatic Preserve** | **27.5837619** | **-82.61594463** | ***Thalassia testudinum*** | **PP430561** |
| **79** | **FL, USA** | **Lower Tampa Bay** | | **1-Aug-22** | **Terra Ceia Aquatic Preserve** | **27.58376283** | **-82.61586358** | ***Thalassia testudinum*** | **PP430562** |
| **80** | **FL, USA** | **Lower Tampa Bay** | | **1-Aug-22** | **Terra Ceia Aquatic Preserve** | **27.58376383** | **-82.61578253** | ***Thalassia testudinum*** | **PP430563** |
| **84** | **FL, USA** | **Lower Tampa Bay** | | **1-Aug-22** | **Terra Ceia Aquatic Preserve** | **27.58368689** | **-82.61618662** | ***Thalassia testudinum*** | **PP430564** |
| **88** | **FL, USA** | **Lower Tampa Bay** | | **1-Aug-22** | **Terra Ceia Aquatic Preserve** | **27.58369066** | **-82.61586249** | ***Thalassia testudinum*** | **PP430565** |
| **93** | **FL, USA** | **Lower Tampa Bay** | | **1-Aug-22** | **Terra Ceia Aquatic Preserve** | **27.58361564** | **-82.61610456** | ***Thalassia testudinum*** | **OR827698** |
| **96** | **FL, USA** | **Lower Tampa Bay** | | **1-Aug-22** | **Terra Ceia Aquatic Preserve** | **27.58361849** | **-82.61586148** | ***Thalassia testudinum*** | **PP430566** |
| **97** | **FL, USA** | **Lower Tampa Bay** | | **1-Aug-22** | **Terra Ceia Aquatic Preserve** | **27.58361941** | **-82.61578043** | ***Thalassia testudinum*** | **OR827705** |
| **101** | **FL, USA** | **Lower Tampa Bay** | | **1-Aug-22** | **Terra Ceia Aquatic Preserve** | **27.58354439** | **-82.61602242** | ***Thalassia testudinum*** | **OR879054** |
| **105** | **FL, USA** | **Lower Tampa Bay** | | **1-Aug-22** | **Terra Ceia Aquatic Preserve** | **27.58354817** | **-82.61569837** | ***Thalassia testudinum*** | **OR827704** |
| **119** | **FL, USA** | **Lower Tampa Bay** | | **1-Aug-22** | **Terra Ceia Aquatic Preserve** | **27.58340475** | **-82.61561522** | ***Thalassia testudinum*** | **PP430548** |
| **128** | **FL, USA** | **Lower Tampa Bay** | | **1-Aug-22** | **Terra Ceia Aquatic Preserve** | **27.58325941** | **-82.6156941** | ***Thalassia testudinum*** | **OR879053** |

| | | | | | | | | |
|---|---|---|---|---|---|---|---|---|
| **132** | **FL, USA** | **Lower Tampa Bay** | **1-Aug-22** | **Terra Ceia Aquatic Preserve** | **27.58318816** | **-82.61561204** | ***Thalassia testudinum*** | **PP430549** |
| **A1** | **FL, USA** | **Dry Tortugas National Park** | **20-May-22** | **Key West** | **24.581857** | **-81.799379** | ***Thalassia testudinum*** | **PP430567** |
| **A2** | **FL, USA** | **Dry Tortugas National Park** | **20-May-22** | **Key West** | **24.581865** | **-81.799295** | ***Thalassia testudinum*** | **PP430568** |
| L1 | FL, USA | Dry Tortugas National Park | 19-May-22 | Marquesas Key | 24.560963 | -82.125589 | *Thalassia testudinum* | |
| **L2** | **FL, USA** | **Dry Tortugas National Park** | **19-May-22** | **Marquesas Key** | **24.560981** | **-82.12555** | ***Thalassia testudinum*** | **PP430569** |
| Q1 | FL, USA | Dry Tortugas National Park | 17-May-22 | Garden Key | 24.626714 | -82.874075 | *Thalassia testudinum* | **PP430570** |
| Q2 | FL, USA | Dry Tortugas National Park | 17-May-22 | Garden Key | 24.626673 | -82.874162 | *Thalassia testudinum* | **PP430571** |
| **R** | **FL, USA** | **Dry Tortugas National Park** | **16-May-22** | **Bush Key** | **24.626357** | **-82.868817** | ***Thalassia testudinum*** | **OR879056** |
| Y | FL, USA | Dry Tortugas National Park | 18-May-22 | Loggerhead Key | 24.635007 | -82.921341 | *Thalassia testudinum* | |
| **Panacea** | **FL, USA** | **Florida Panhandle** | **10-May-23** | **Panacea** | **29.99911** | **-84.35056** | ***Thalassia testudinum*** | **OR879055** |
| MAIRIM | Puerto Rico | Jobos Bay National Estuarine Research Reserve | 1-Feb-23 | MAIRIM Site | Not available | Not available | *Halophila stipulacea* | |
| CPL | Puerto Rico | Jobos Bay National Estuarine Research Reserve | 1-Feb-23 | Cayo Puerca Lulu | Not available | Not available | *Halophila stipulacea* | |
| Z1 | VA, USA | York River | 2-Aug-17 | York River State Park | Not available | Not available | *Zostera marina* | |
| Z2 | MA, USA | West Falmouth Harbor | Not available | West Falmouth Harbor | Not available | Not available | *Zostera marina* | |
| Z3 | AK, USA | Sitka | Not available | Sitka | Not available | Not available | *Zostera marina* | |
| Z4 | New Zealand | Not available | Dec-22 | New Zealand | Not available | Not available | *Zostera marina* | |
| Z5 | Kalmar, Sweden | Not available | Not available | Kalmar | Not available | Not available | *Zostera marina* | |
